# The perceived position of a moving object is reset by temporal, not spatial limits

**DOI:** 10.1101/2021.12.14.472615

**Authors:** Sirui Liu, Peter U. Tse, Patrick Cavanagh

## Abstract

When the internal texture of a Gabor patch drifts orthogonally to its physical path, its perceived motion deviates dramatically from its physical path. The local position shifts accumulate to such an extent that a 45° oblique physical path appears to be vertical. However, at some point, a limit is reached and the path resets back to its veridical location, whereupon a new accumulation starts, making the new perceived path segment appear parallel to the pre-reset segment, but offset horizontally from it. Here, we tested whether spontaneous resets of this motion-induced position shift depend on the time or the distance over which position errors accrue, or both. We introduced a temporal gap in the middle of the path that forced the illusory path to reset back to its veridical physical position. This gap-triggered reset allowed us to measure the magnitude of the illusory offset up to that point. We found that perceived offset was less than expected for the angle of illusory drift, indicating that spontaneous resets had occurred prior to the gap-induced reset. The position offset decreased when the pre-gap duration increased but approximately doubled when the path length doubled. This pattern of perceived offsets is best accounted for by spontaneous resets that occur randomly over time at a constant rate, independently of the distance traveled. Our results suggest a temporal, not spatial, limit for the accumulation of position errors that underlies this illusion.

## Introduction

A moving Gabor patch with orthogonally drifting internal texture is perceived to deviate dramatically from its physical path. Compared with other well-known motion-induced position shift effects, this ‘double-drift’ stimulus (Lisi & Cavanagh, 2015; also called ‘the curveball illusion’, Kwon, Tadin, & Knill, 2015, Shapiro, Lu, Huang, Knight, & Ennis, 2010; or ‘infinite regress illusion’, Tse & Hsieh, 2006) has a much a larger perceived position displacement that can deviate from its physical location by several degrees of visual angle (**Figure 1**). One distinctive feature that makes the double-drift stimulus so powerful is that its perceived motion path is formed by integrating motion signals and accumulating position shifts orthogonal to the veridical path over a second or more, while other motion-induced position shift stimuli, such as the flash-grab (Cavanagh & Anstis, 2013) and motion drag (De Valois & De Valois, 1991) stimuli, only integrate motion signals over 80 to 100 ms (Cavanagh & Anstis, 2013; Chung et al., 2007; Kosovicheva et al, 2014; Joen et al, 2020). Lisi and Cavanagh (2015) demonstrated that the illusion is not just a change in perceived direction but affects perceived position as well. In particular, they showed that a temporal gap of 250ms in the middle of the path resets the position offset such that the post-gap segment appears to start from its new physical location rather than from its previous perceived location (**Figure 2**). They argued that if this illusion only involved a change of direction without affecting perceived position, the stimulus would appear at the same location after the temporal gap (Lisi & Cavanagh, 2015). Kwon, Tadin, and Knill (2015) presented a computational model according to which position and motion information are estimated based on past sensory signals when the precision of current position information is low. They found that the illusion got stronger the further in the periphery it was placed, and that the perceived speed of the Gabor’s internal motion slowed, suggesting that its energy was being captured to create the illusory motion direction. Indeed, the illusion is strongest on an equiluminant background in the periphery where position uncertainty is highest (Cavanagh & Tse, 2019). In this case, the target’s motion starts to contribute to its position estimate, generating displacements in the direction of the target’s motion, predicting where it should be next. However, the target’s motion direction is taken to be a combination of both the within-target texture motion and its translational motion, sending the estimates of the new position off in the illusory, combined direction. When the same Gabor is placed on a dark background, position uncertainty is low. In this case, internal motion does not contribute to new position estimates and there is no illusion (Cavanagh & Tse, 2019).

**Figure 1.**
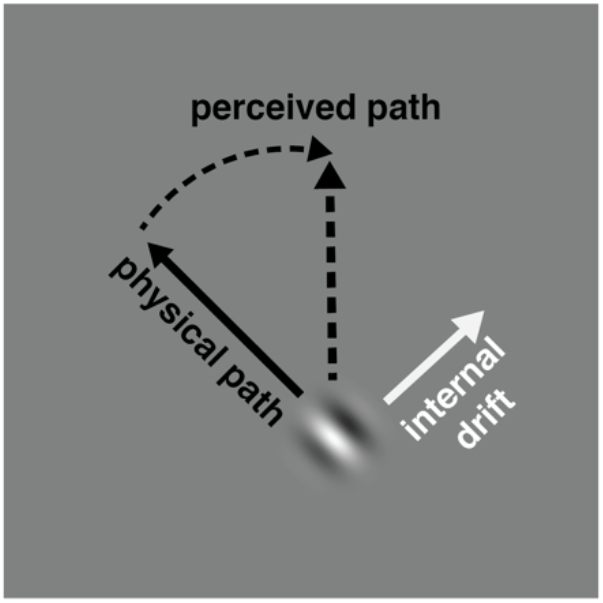
Double-drift stimulus. A Gabor patch that is moving obliquely (physical path) can be perceived to be moving vertically (perceived path) if its internal texture drifts orthogonally to the physical path (up and to the right).

**Figure 2.**
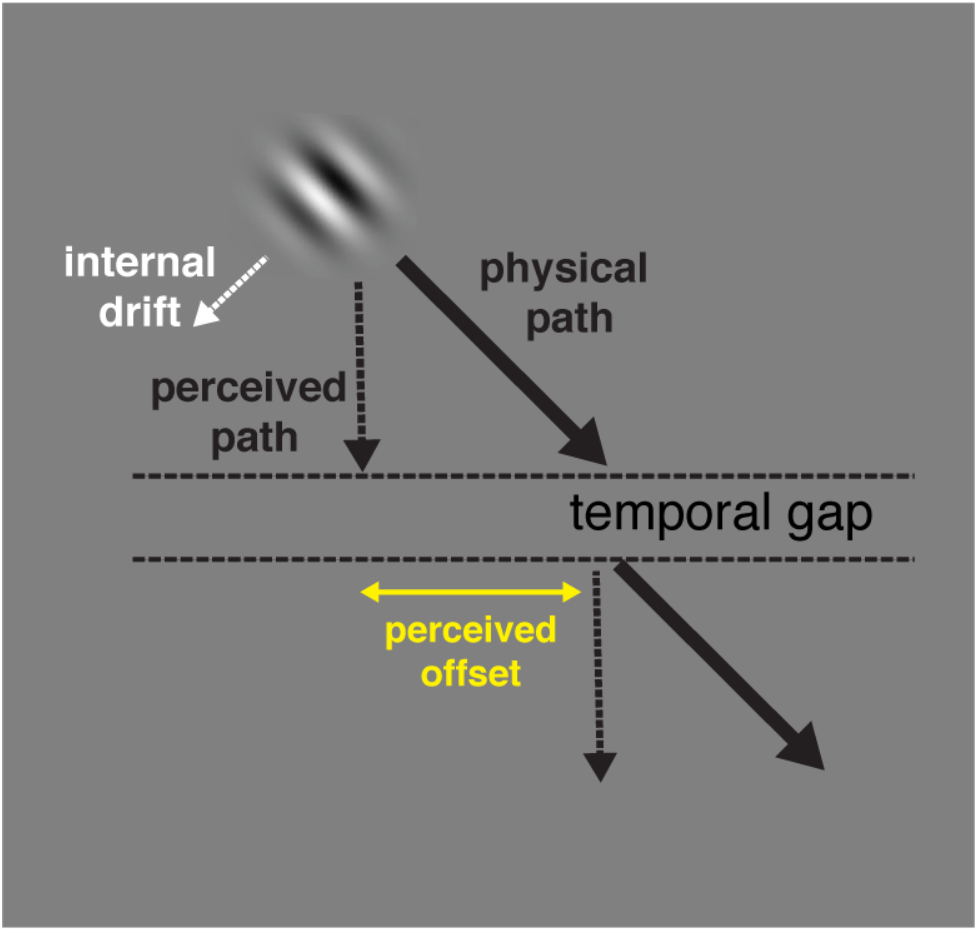
Schematic demonstration of a position reset for a double-drift stimulus caused by a temporal gap. A Gabor patch moves downward 45° to the right with its internal motion orthogonal to its motion direction such that the direction of its motion path appears to be vertical. A blank temporal gap is introduced in the middle of the path and the Gabor reappears at the same physical location as before the gap and continues its movement in the same direction. Rather than starting from its last perceived pre-gap location, the Gabor appears to be shifted to the new physical location and starts to accumulate position errors from that location, creating a perceived offset between the two motion segments.

In this study, we further explored the nature of the motion-position integration process that underlies the illusion. Kwon et al. (2015) and Shapiro et al. (2010) reported that the illusory shift saturates into a curved trajectory as the path length increases. In contrast, we have frequently observed something quite different – the path continues linearly first and then at some point, a limit is reached. When that happens, the path resets back toward its physical location and a new linear accumulation starts. These spontaneous resets appear to be similar to the forced reset created by a temporal gap used by Lisi and Cavanagh (2015). In both cases, a new linear, illusory motion segment begins parallel to the first, starting from the physical location (**Figure 3**). The consequence of these resets is that the whole path appears to saturate into a curved trajectory with averaging over trials, so that the existence of the individual resets from a linear path is masked. In a separate study, Nakayama and Holcombe (2020) asked participants to trace or draw the motion paths they saw for the double drift stimuli and the presence of the resets was clear in this case, triggered by distracting flashes on the screen (**Figure 4**).

**Figure 3.**
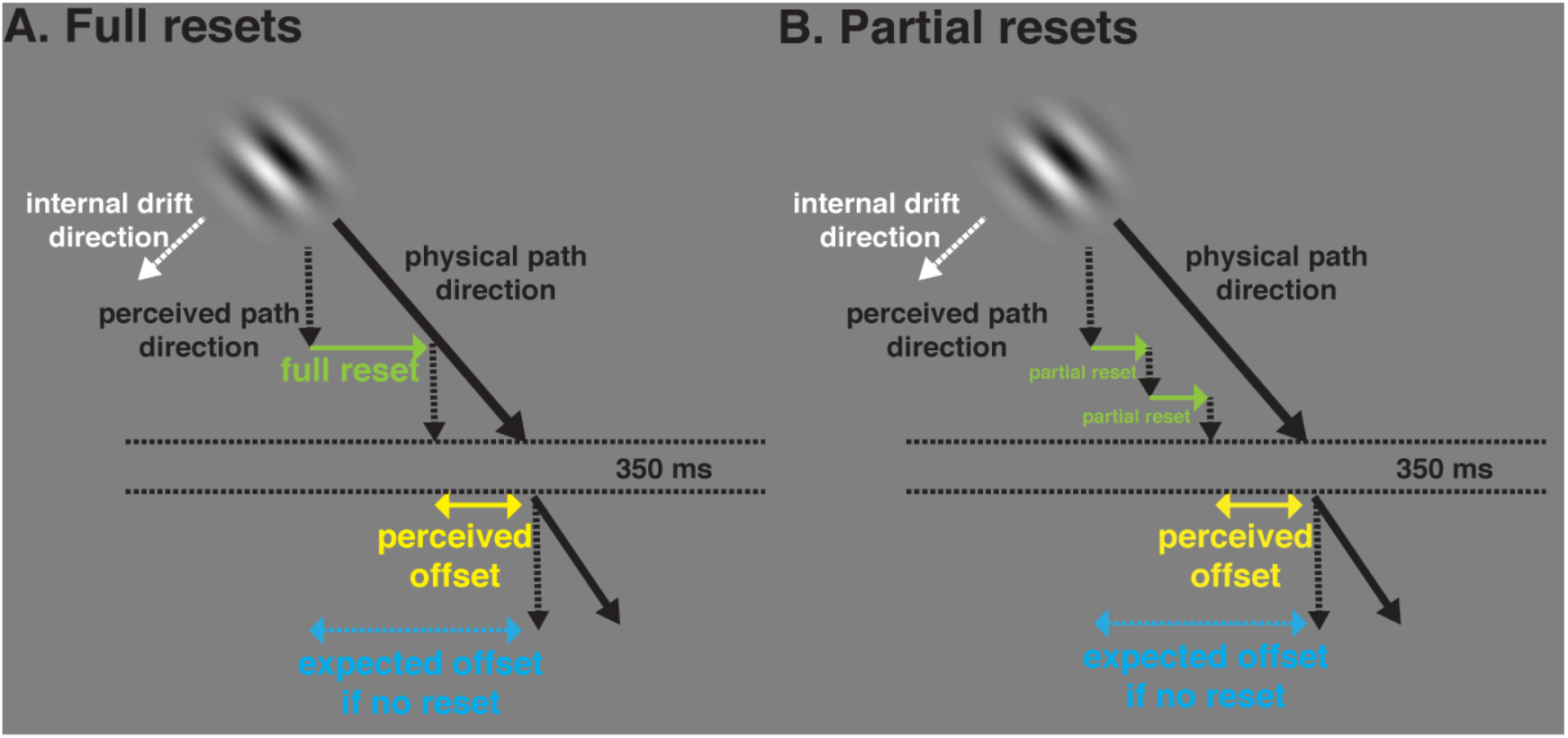
Schematic demonstration of a spontaneous position reset for a double-drift stimulus. A Gabor patch moves downward 45° to the right with its internal motion orthogonal to its motion direction such that the direction of its motion path appears to be vertical. A blank temporal gap (350ms) is introduced during the movement and the Gabor reappears at the same physical location as before the gap and continues its movement in the same direction. If there is a spontaneous position reset before the temporal gap (green arrow), the path resets back toward its physical location and a new accumulation starts, making a new perceived vertical segment parallel to the first but offset horizontally. The perceived position may either reset fully (A) or partially (B) back to the physical location. If the accumulation has been linear and free of loss before the temporal gap, the expected offset size (blue arrow) can be calculated from the angle of the illusory deviation. If the perceived offset (yellow arrow) is less than this expected offset size, it indicates that some position reset or saturation has occurred before the temporal gap.

**Figure 4.**
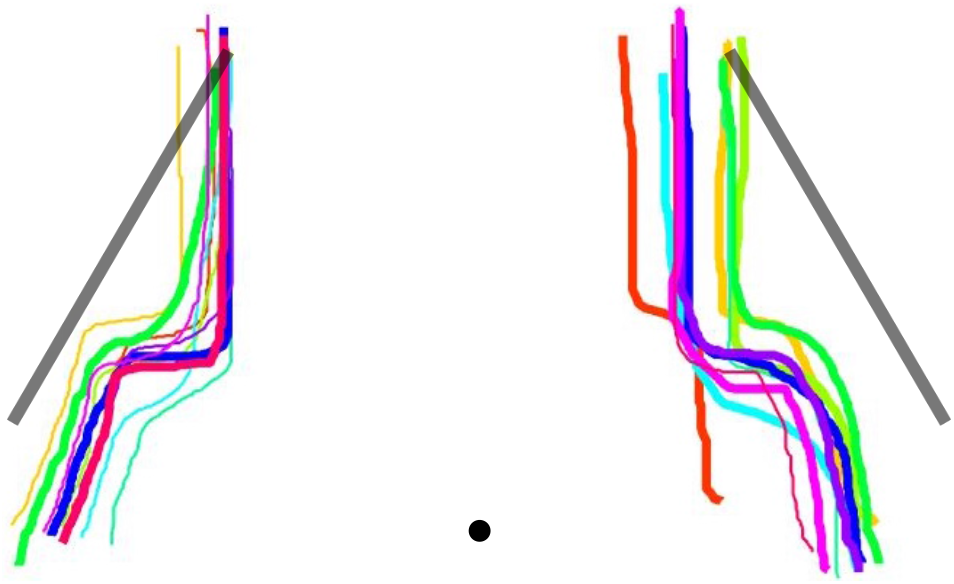
Individual traces of the double-drift path. Two Gabor patches moved downward obliquely (black line) with their internal drifts driving the perceived paths vertical. Flashes were presented on both sides of the stimuli part way through their motion trajectories. Participants observed a single transit of motion and then traced their impression of the paths using the computer mouse, each showing a clear reset triggered by the flashes. From Nakayama and Holcombe (2020).

The source of spontaneous position resets has not yet been examined. Here we investigated whether the upper limit for the accumulation of the illusory position shift depends on the time or/and the distance traveled by the double-drift stimulus. Specifically, if a reset is triggered once the illusory offset exceeds a maximum distance from the Gabor’s real position, regardless of the time of accumulation, it suggests that there is a critical spatial error, or a ‘red line’, for the illusory offset. Conversely, if the reset is triggered solely on the basis of the time since the illusory offset began, it suggests a temporal limit on the accumulation of the position displacements. If this temporal limit is in the range of seconds, it suggests that the accumulation process must logically involve higher-order brain regions, as cells in early processing stages do not have integration time windows on the scale of seconds; these are found instead in frontal areas that support visual working memory maintenance (Funahashi, Bruce, & Goldman-Rakic, 1989; 1990). Indeed, a recent fMRI study using multivariate pattern analysis found that the perceived path of the double-drift stimulus did not share activity patterns with a matched physical path in early visual areas traditionally associated with visual processing. This shared representation was only found in anterior brain regions such as the frontal cortex (Liu, Yu, Tse, & Cavanagh, 2019). Lastly, both distance and time may contribute to determine the moment of the reset. For example, the longer the perceived position is beyond the “red line” for error, the more likely the reset. There are a number of ways that space or time or both could influence the occurrence of resets. Here we will try to evaluate only a few simple possibilities.

How can we measure these resets? Since a temporal gap in the path can force a reset, as shown in Lisi and Cavanagh (2015), the perceived offset between pre- and post-gap motion path should reveal the magnitude of the accumulation up to the reset. The present study thus measured how much position error has accumulated up to a certain point by forcing a reset with a temporal gap at that point. Specifically, we presented a double-drift stimulus over different time durations and path lengths before the temporal gap, followed by a similar continued motion after the gap. We compared the perceived offsets between the two path segments with the expected size of offset that would be seen if no spontaneous reset(s) had occurred before the gap. Critically, to allow for a prediction of what the perceived offset would be without resets, we asked each participant to first adjust the speed of the internal motion so that the perceived path angle (ignoring any position resets that they saw) was always, locally, 45°. If the accumulation is linear and free of loss during the first segment, the offset of the perceived location from the physical location must be 70.7% (or 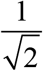) of the physical path length for a 45° angle of the illusory deviation. Anything less than 70.7% (for this 45° illusion size) indicates that some spontaneous reset or saturation must have occurred before the temporal gap (**Figure 3**). Overall, we found that the perceived offset was smaller than expected and decreased for longer durations with the same path length, but approximately doubled when the path length was doubled. The observed data were best explained by partial resets that happen randomly over time with a fixed probability, regardless of the illusory offset from the physical path.

## Methods

### Participants

Twenty students from Dartmouth College participated (Ten in the main experiment: 7 females; age range: 18–38, mean age = 23.4 ± 6.8; Ten in the control experiment: 8 females; age range: 18–20, mean age = 18.9 ± 0.7) and all reported normal or corrected-to-normal vision. All volunteered to participate and were naïve to the purpose of the study. Participants signed an informed consent approved by the Institutional Review Board at Dartmouth College and received a compensation of course credits or $10/hour.

### Apparatus

Stimuli were generated on an Apple iMac Intel Core i5 (Cupertino, CA) using MATLAB R2015a (MathWorks, Natick, MA) and PsychToolbox-3 (Kleiner et al., 2007) and were displayed in a dark room on a 16” ViewSonic G73f CRT monitor (1024 768 pixels at 60 Hz) placed 57 cm from the observer. The participant’s head was stabilized using a chinrest during the psychophysical experiment.

### Stimuli

A fixation point (0.2 degrees of visual angle [dva] in diameter) was presented at 5 dva to the left of the screen throughout the experiment. The stimulus was a Gabor patch (a sinusoidal grating within a Gaussian envelope) with a spatial frequency of 1 cycle/dva and 100% Michelson contrast (Peli, 1990). The standard deviation of the Gaussian envelope was 0.15 dva. The Gabor patch moved along a linear path at 45° either leftward or rightward from vertical. The midpoint of the trajectory was placed at 5 dva to the right of the screen center so that the stimulus was 10 dva away from fixation. The sinusoidal grating had the same orientation as the motion path. Stimuli were presented on a uniform grey background with luminance of 53 cd/m^2^, equal to the mean luminance of the Gabor. There were two parts of the experiment. In the first part, the Gabor patch traversed back and forth along a linear path (external motion). The internal texture of the Gabor also drifted in the orthogonal and downward direction of the external motion path (internal motion) to produce a perceived vertical motion path. The internal motion reversed its direction each time the external motion reversed. The speed of the internal motion was first presented at a random frequency that ranged from 0.5 to 5.5 Hz on each trial and was adjusted, as described below, to make the local angle of the Gabor path appear vertical for each path length, duration and direction of internal motion. In the second part of the experiment, the same stimulus was used; however, it was only presented for half a cycle on each trial, and always started moving downward from the top position. The internal texture either drifted (‘double-drift stimulus’) or stayed static (‘control stimulus’). When the Gabor reached the midpoint of the path, there was a brief temporal gap of 350ms with the stimulus removed from the screen. After the blank gap, the stimulus would reappear at the same height but at a shifted horizontal position (−3.5, −2.1, −0.7, 0.7, 2.1, 3.5 dva) and continued its movement in the same direction for the same amount of time and path length as before the gap. Three durations (1s, 2s, 3s) and two path lengths (2 dva, 4 dva) before the gap were used in both parts of the experiment (**Figure 5**).

**Figure 5.**
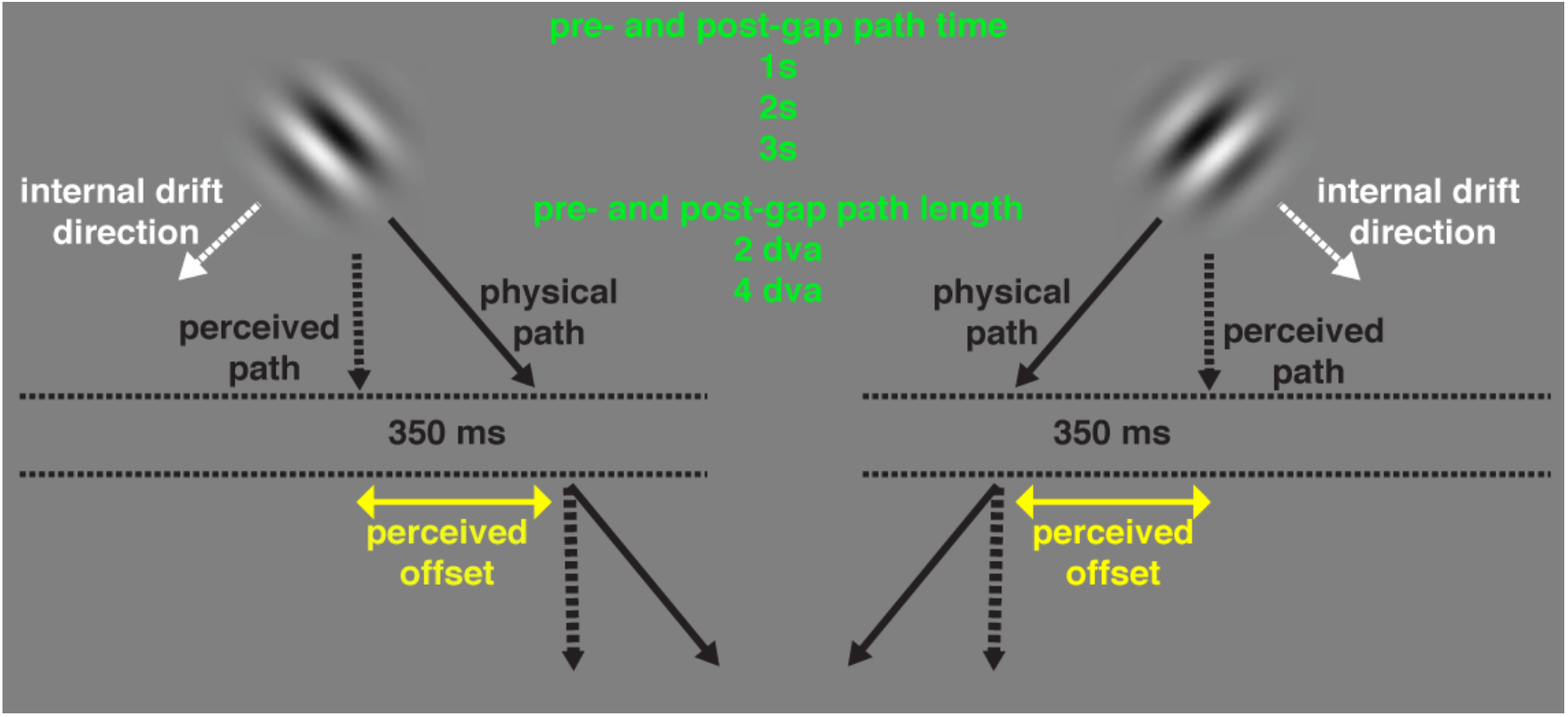
Stimulus demonstration and experiment conditions. A Gabor patch moved along a linear path downward at 45° to the right or left from vertical. Its internal texture also drifted in the orthogonal and downward direction of the physical path to produce a vertical perceived motion path. A blank temporal gap (350ms) was introduced during the movement and the Gabor reappeared at the same physical location as before the gap and continued its movement in the same direction for the same amount of time and path length as before the gap. The physical motion path had two possible orientations (leftward or rightward from vertical). The path before and after the gap had three possible durations (1s, 2s, 3s) and two possible path lengths (2 dva and 4 dva). If a spontaneous position reset occurs during the blank gap, the post-gap motion segment should appear parallel to the pre-gap segment but offset horizontally by less than the full amount expected.

### Procedure

Participants were instructed to fixate at the fixation point throughout all experiments. Each participant completed two separate parts of the experiment. In the first part, a Gabor patch was shown in the periphery and moved back and forth along a linear path with its internal texture drifting orthogonally relative to the external motion. In each trial, participants adjusted the speed of the internal drift by pressing the up arrow key to increase the speed and the down arrow key to decrease the speed until the physical trajectory appeared locally vertical along its path, ignoring any sidesteps from resets. The purpose of this task was to find the internal motion speed that produced a perception of locally vertical motion for physical paths either 45° leftward or rightward from vertical at each of the different external motion speeds used in the subsequent task. This adjustment produces a constant perceived local path angle in all conditions. This allows us to predict the expected amount of shift throughout so that the differences in perceived offset between the two motion segments are not simply due to differences in the perceived path angle. Each participant completed 15 adjustment trials for each external motion path orientation, stimulus duration, and path length for a total of 180 trials divided in 2 blocks. The average of the internal motion speed for each stimulus condition was calculated individually and was then used for that participant in the following task for that same condition. See **Figure 6** for group averaged internal motion speeds for each condition as well as the expected angle of the illusory path derived from the vector sum of the external motion speed and the adjusted internal motion speed, a model that was found to produce the perceived motion direction when position uncertainty is high (Cavanagh and Tse, 2019). In the second part of the experiment, in each trial, after a 400 ms fixation period, the same Gabor patch as in the first task was shown in the periphery traversing its path once with a temporal gap of 350 ms presented in the middle of the path during which nothing was presented. After the stimulus reached the final position of the motion path it was removed from the screen and participants reported the direction of the horizontal position shift between the first and second half of the path by pressing a key corresponding to left or right shift. Each participant completed 20 trials for each external motion path orientation, size of horizontal position shift, stimulus duration, and path length for a total of 1440 trials that were divided in 4 sessions on separate days, each composed of 6 blocks of 60 trials with all conditions randomized and counterbalanced across trials. A control stimulus with no internal motion was used in a different group of participants as a comparison.

**Figure 6.**
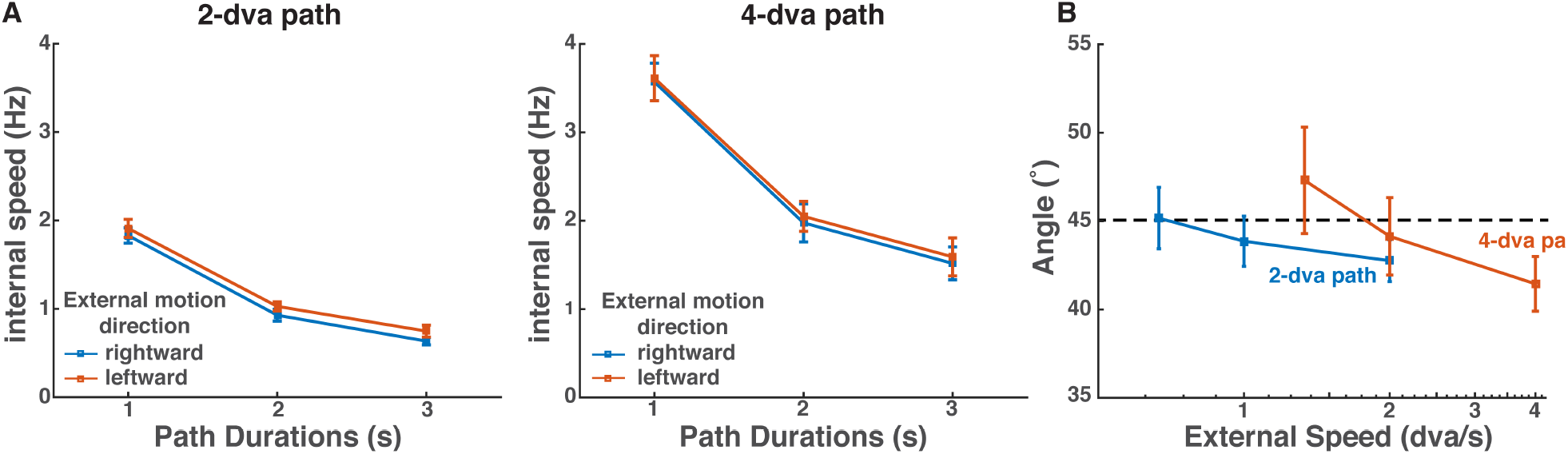
Mean internal motion speeds across participants (n =10). **A.** Group-averaged internal motion speeds that produced a perception of vertical motion for each time duration and path length conditions of the leftward and rightward moving double-drift stimulus in the adjustment task. Error bars represent ±1 *SEM*. Stimuli with higher external motion speed (i.e. shorter path duration and longer path length) need significantly higher internal motion speed to produce the same perceived path angle as those with lower external motion speeds. **B.** Illusory path orientation 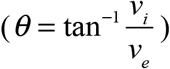 derived from the vector sum of the adjusted internal motion speed (*v_i_*) and external motion speed (*v_e_*) as a function of the external speed. The dashed line represents the path angle (45°) that the participants were instructed to adjust to.

### Analysis

#### PSE estimation

The percentage of trials with rightward responses was calculated as a function of the size of the physical horizontal position shift of the stimulus after the temporal gap. The data were fitted using a cumulative Gaussian psychometric function:

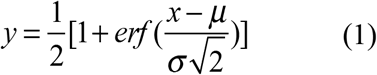

in which *y* is the proportion of trials of rightward responses; *x* is the size of physical offset between the two motion segments; *μ* is the location parameter; and *σ* is the scale parameter, using a maximum likelihood procedure. The point of subjective equality (PSE) – the point at which the two motion segments were subjectively aligned so that rightward and leftward were reported equally – was interpolated from individual function fits for each condition. The average of these PSE values for each condition across participants was then calculated. To test whether and how time duration and path length affected the magnitude of position shift up to the temporal gap, we combined the PSEs from the two external motion conditions after flipping the signs of the PSEs from the rightward moving condition and conducted repeated-measure ANOVAs and post hoc pairwise *t* tests with multiple tests correction using the false discovery rate (FDR; Benjamini and Hochberg 1995) on these perceived offset values.

#### Model fitting

We explored models that had spatial or temporal limits that would trigger a reset. The resets could either reduce the offset from the physical location to 0 (‘full resets’) or reduce the offset by a portion of the distance to last reset (‘partial resets’). In all cases, we assume that rate of accumulation of the illusory offset is linear (see **Figure 3**) and determined by the external speed of the Gabor (path length, *P*, divided by pre-gap duration, *T*) and the local angle, *α*, of the illusory path (which is titrated by changing the internal speed to be always 45°), such that the magnitude of illusory offset, *S*, at time point *t* since last reset is given by:

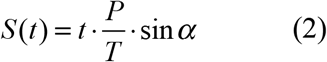

We compared the perceived offset values for each stimulus condition with predictions from two types of models: 1) a fixed time limit model or fixed space limit model where a reset happens after a fixed time interval or a fixed distance that the illusory path has drifted away from the physical path; 2) a random time limit or random space limit where a reset happens randomly with a probability per second or per dva that the stimulus has drifted away from its physical path. For each path length and duration, we simulated the experiment 10,000 times for each possible parameter value of each model, calculated the average of the drift values across runs at each time point and then calculated the root mean squared error (RMSE) of the predicted drift values for that condition. For each best fit model, we calculated the Akaike Information Criterion (*AIC*), that is given by 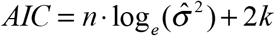, in which *n* is the sample size, 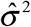 is 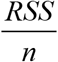 (*RSS* = residual sum of squares; sum of the deviations between the data and the estimation of that model), and *k* is the number of parameters. We then corrected these *AIC* values for small sample size, 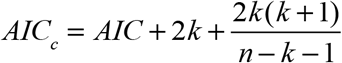, as suggested by Burnham and Anderson (2003), as well as their difference relative to the model with the lowest *AIC_c_*, Δ*AIC_c_*, to compare model performance.

## Results

**Figure 7** show the group-averaged percentage of trials on which participants perceived the post-gap motion trace to be to the right of the pre-gap motion trace of the double-drift stimulus as a function of the presented physical offset between motion traces. Psychometric curves were fitted individually (as given in Equation 1) to estimate the PSEs where the pre- and post-gap motion segments appeared aligned. PSEs from these psychometric curves indicate the physical shift needed to null the perceived offset of the two motion segments between the temporal gap. We found that for all time and path length conditions, the curves for the rightward moving double-drift stimulus was shifted to the left relative to the condition when the Gabor had no internal drift (**Figure 7 and 8** left panel). This indicates that when a double-drift stimulus physically moving obliquely to the right reappeared in the same physical location after the temporal gap, observers were more likely to perceive the stimulus to be shifted rightward from where the first segment ended. In other words, the temporal gap had reset the path to the right of the perceived end point of the first motion segment. Thus, a horizontal position shift to the left of the post-gap motion path would null the perceived offset produced by the gap-induced position reset. Conversely, when the double-drift stimulus was moving obliquely to the left, the curve was shifted to the right relative to the control condition (**Figure 7 and 8** right panel), indicating that the post-gap segment was more likely to be perceived as shifted leftward back to its physical position after the temporal gap. Results were highly similar across all observers and the observed bias in PSEs in the opposite direction of the external motion was only observed for the double-drift stimulus [*F*_(1,9)_ = 78.31, *p* < 0.001, *η*^2^ = 0.70] but not for the control stimulus with no internal motion [*F*_(1,9)_ = 1.72, *p* = 0.22, *η*^2^ = 0.07]. This pattern of results is consistent with those found in Lisi and Cavanagh (2015) where they showed that participants would judge the position after the temporal gap relative to the illusory position that was built up at the end of the first half of the motion path, suggesting that the illusion involves an accumulation of position displacements.

**Figure 7.**
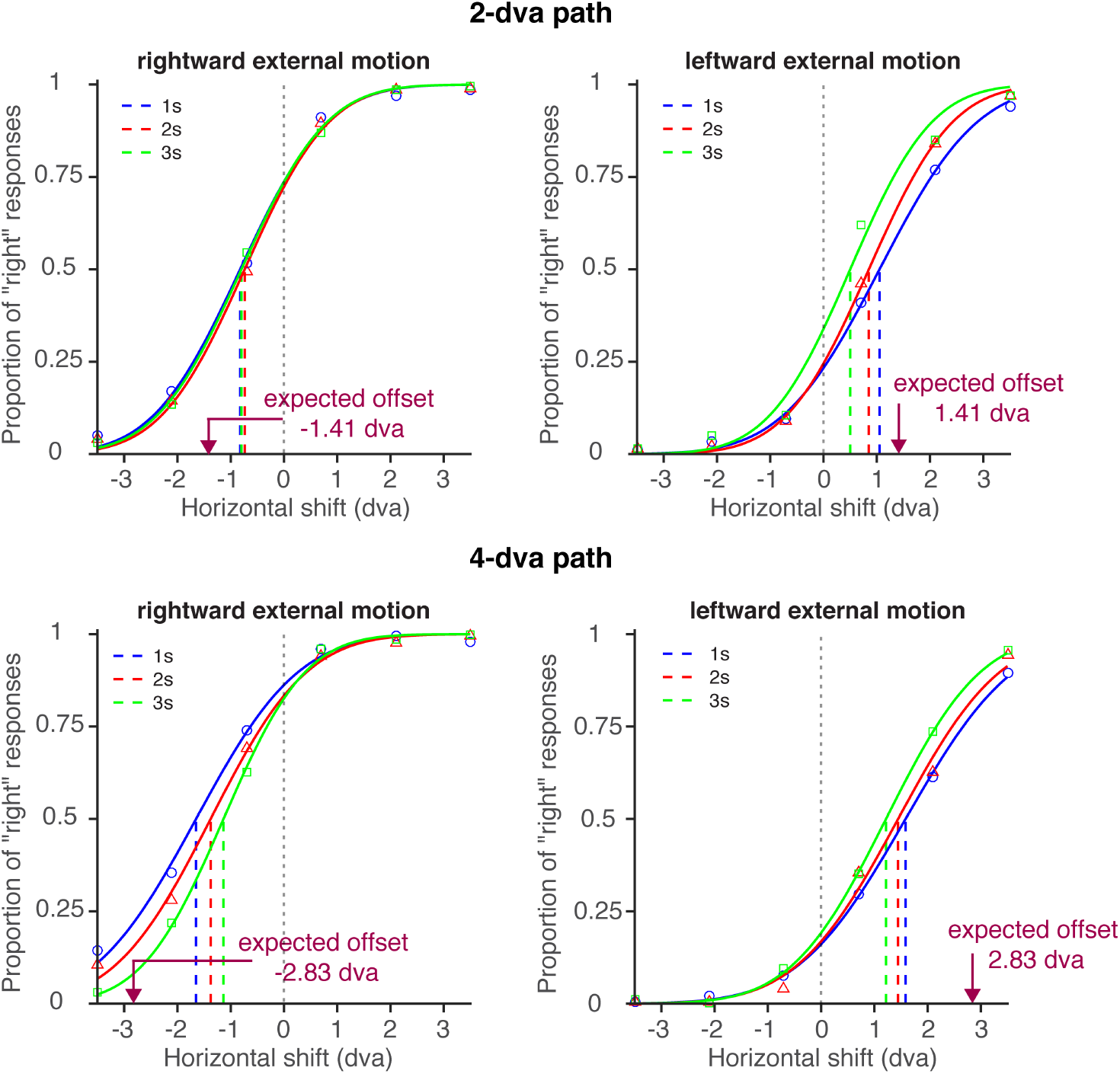
Psychometric functions of position judgment for the double-drift stimulus. Group-averaged percentage of trials on which participants perceived the Gabor to be shifted rightward after the temporal gap as a function of the physical offset between the motion traces for the rightward and leftward moving double-drift stimulus. The point of subjective equality (PSE) of each curve represents the physical offset that yielded 50% right responses for that stimulus condition.

**Figure 8.**
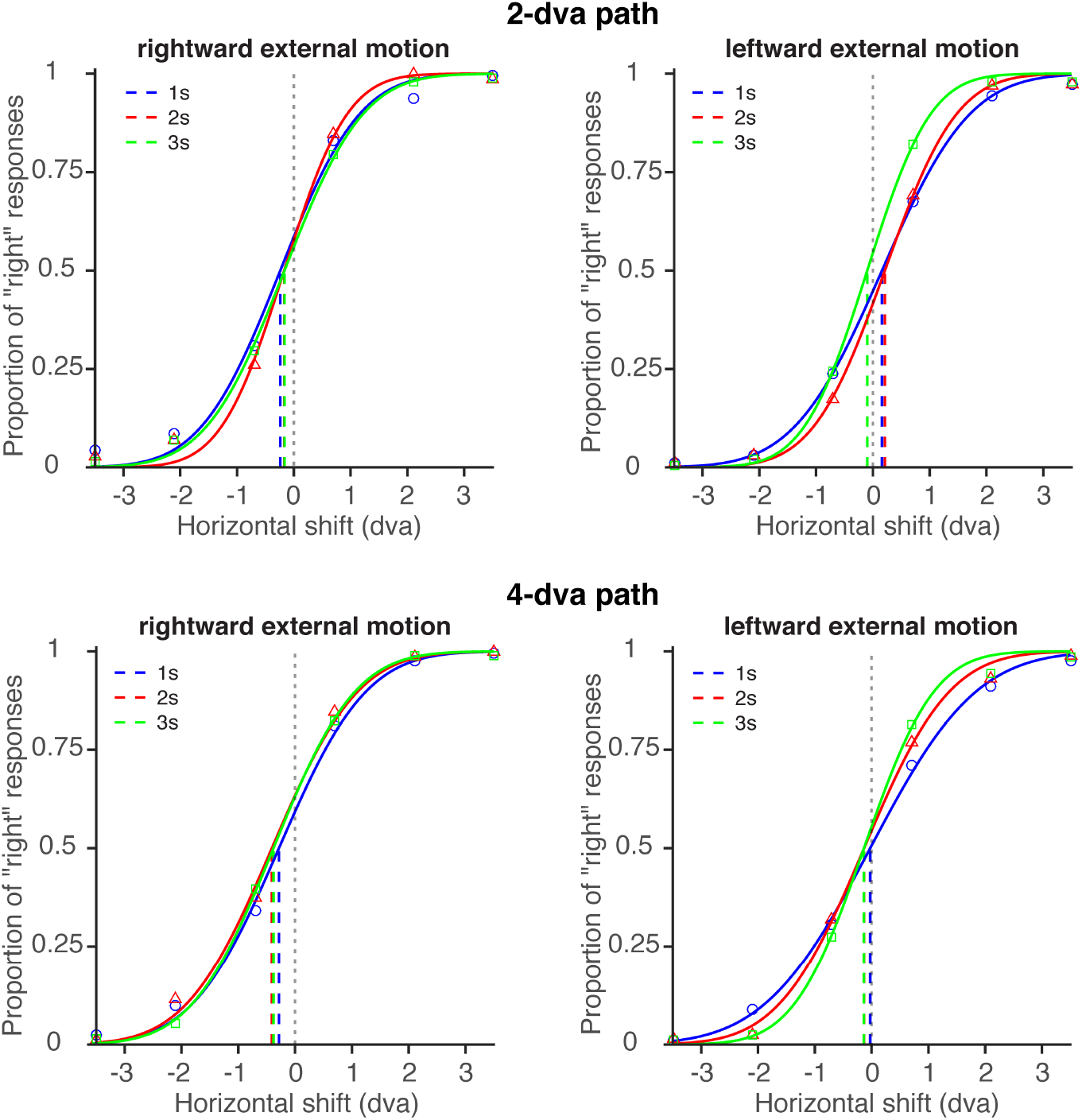
Psychometric functions of position judgment for the control stimulus. Group-averaged percentage of trials on which participants perceived the Gabor to be shifted rightward after the temporal gap as a function of the physical offset between the motion traces for the rightward and leftward moving Gabor with no internal motion (control stimulus). The point of subjective equality (PSE) of each curve represents the physical offset that yielded 50% right responses for that stimulus condition.

Next, we analyzed the effect of stimulus duration and path length on the PSEs for the leftward and rightward moving double-drift stimulus. Results showed that there was a significant main effect of time duration (rightward external motion: *F*_(2,18)_ = 3.95, *p* = 0.05, *η^2^* = 0.03; leftward external motion: *F*_(2,18)_ = 6.58, *p* = 0.02, *η*^2^ = 0.07) and path length for both external motion directions (rightward external motion: *F*_(1,9)_ = 9.79, *p* = 0.01, *η*^2^ = 0.18; leftward external motion: *F*_(1,9)_ = 11.96, *p* = 0.007, *η*^2^ = 0.18), and a significant interaction between the two for the rightward [*F*_(2,18)_ = 6.66, *p* = 0.01, *η*^2^ = 0.02] but not for the leftward moving double-drift stimulus [*F*_(2,18)_ = 1.15, *p* = 0.34, *η*^2^ = 0.005]. Since we found no significant difference in the magnitude of perceived offset between the leftward and rightward moving double-drift stimulus for each path length and stimulus duration condition (all FDR-adjust *p*s > 0.1), we combined the data from the two external motion conditions after flipping the signs of the PSEs from the rightward moving condition (**Figure 9**). There was a significant effect of time on PSEs in the 2-dva path length condition [*F*_(2,18)_ = 9.47, *p* = 0.006, *η*^2^ = 0.11], reflecting significant differences among the three time conditions, with longer time durations leading to smaller perceived offsets [1s vs. 2s: *t*_(9)_ = 2.5, *p* = 0.03, Cohen’s d = 0.42; 2s vs. 3s: *t*_(9)_ = 2.85, *p* = 0.03, Cohen’s d = 0.41; 1s vs. 3s: *t*_(9)_ = 3.39, *p* = 0.02, Cohen’s d = 0.84]. Similarly, for the 4-dva path length condition, a significant effect of time on PSEs was observed [*F*_(2,18)_ = 5.86, *p* = 0.02, *η*^2^ = 0.11; 1s vs. 2s: *t*_(9)_ = 1.04, *p* = 0.33, Cohen’s d = 0.21; 2s vs. 3s: *t*_(9)_ = 3.12, *p* = 0.04, Cohen’s d = 0.62; 1s vs. 3s: *t*_(9)_ = 2.69, *p* = 0.04, Cohen’s d = 0.76]. Pairwise comparisons between the two path lengths for each time condition showed a significant increase in perceived offset for longer distance for each stimulus duration condition [1s: *t*_(9)_ = 4.19, *p* = 0.002, Cohen’s d = 1.13; 2s: *t*_(9)_ = 4.37, *p* = 0.002, Cohen’s d = 1.44; 3s: *t*_(9)_ = 4.16, *p* = 0.002, Cohen’s d = 1.28]. Together, these results showed that the perceived offset was smaller than expected for all tested path lengths and durations, decreased for longer durations with the same path length, but approximately doubled when the path length was doubled.

**Figure 9.**
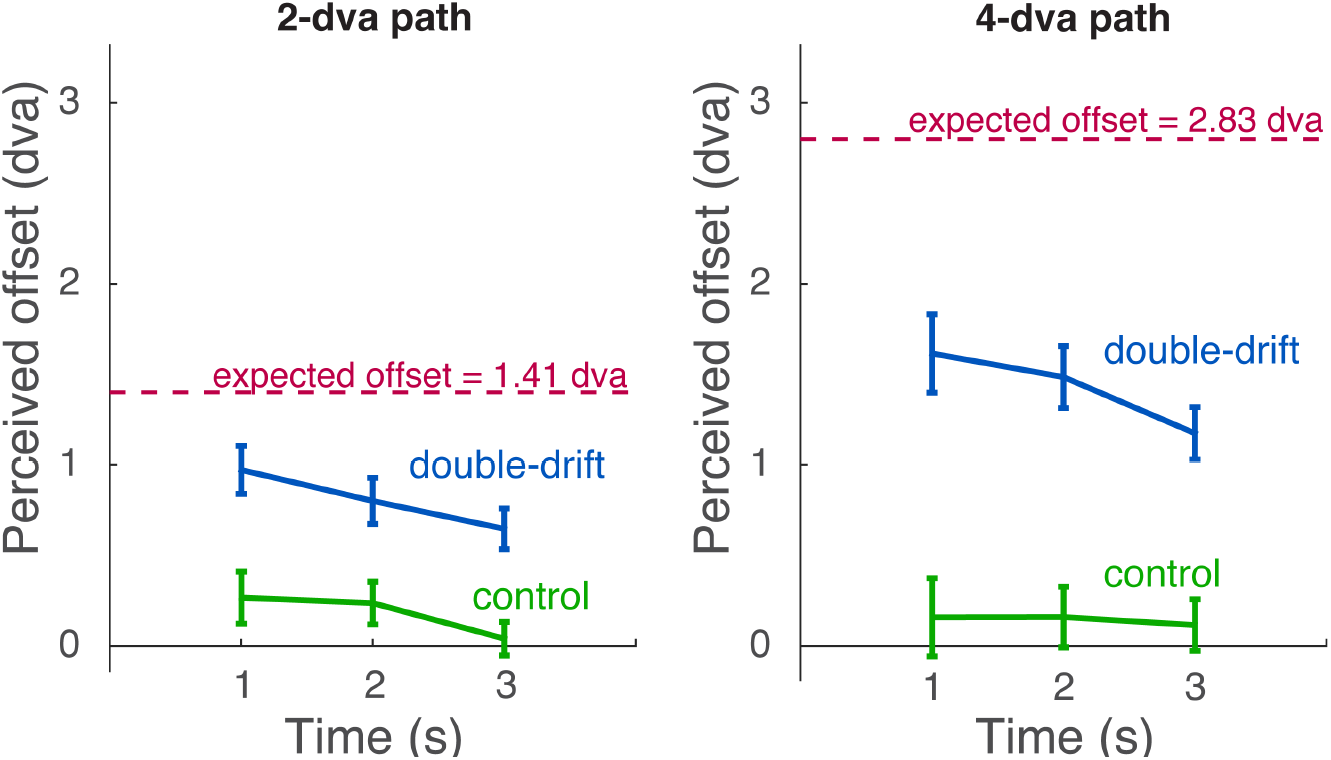
Perceived offset (n = 10). Group-averaged perceived offset values of the double-drift and control stimulus after combining the data from the two external motion conditions by flipping the signs of the PSEs from the rightward moving condition for each time and path length conditions. Red horizontal lines show expected PSEs for each path length if there were no spontaneous resets of the double-drift stimulus. Error bars represent ± 1 *SEM*.

Lastly, we examined models with spatial or temporal limits that would trigger a reset and compared the predictions of each model with the observed perceived offset values. **Figure 10** and **11** show examples of the best fitting averaged drift values across 10,000 repetitions of simulation of each model. **Table 1** shows the average number of resets across runs of simulation per condition for each best fitting model along with model performance.

**Figure 10.**
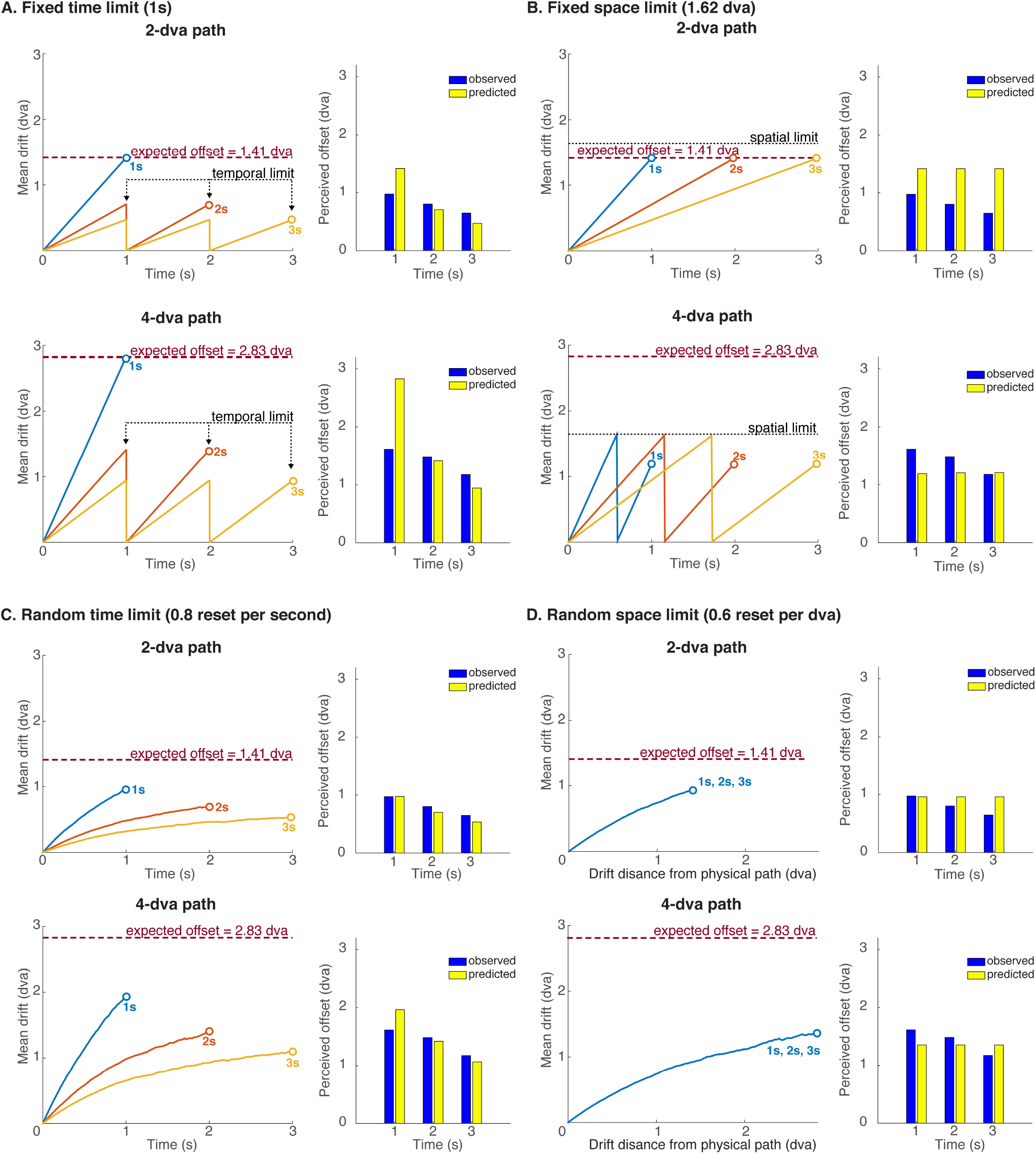
Full reset model comparison. Average drift values across 10,000 simulations at each time point using the parameters that yielded lowest root mean square error (RMSE) between the observed PSEs and the predicted values (left panel) and the predicted and observed values at each path duration (1s, 2s and 3s) (right panel) for the A) fixed time limit model at 1s, B) fixed space limit model at 1.62 dva, C) random time limit model at 0.8/s and D) random space limit model at 0.6/dva.

**Figure 11.**
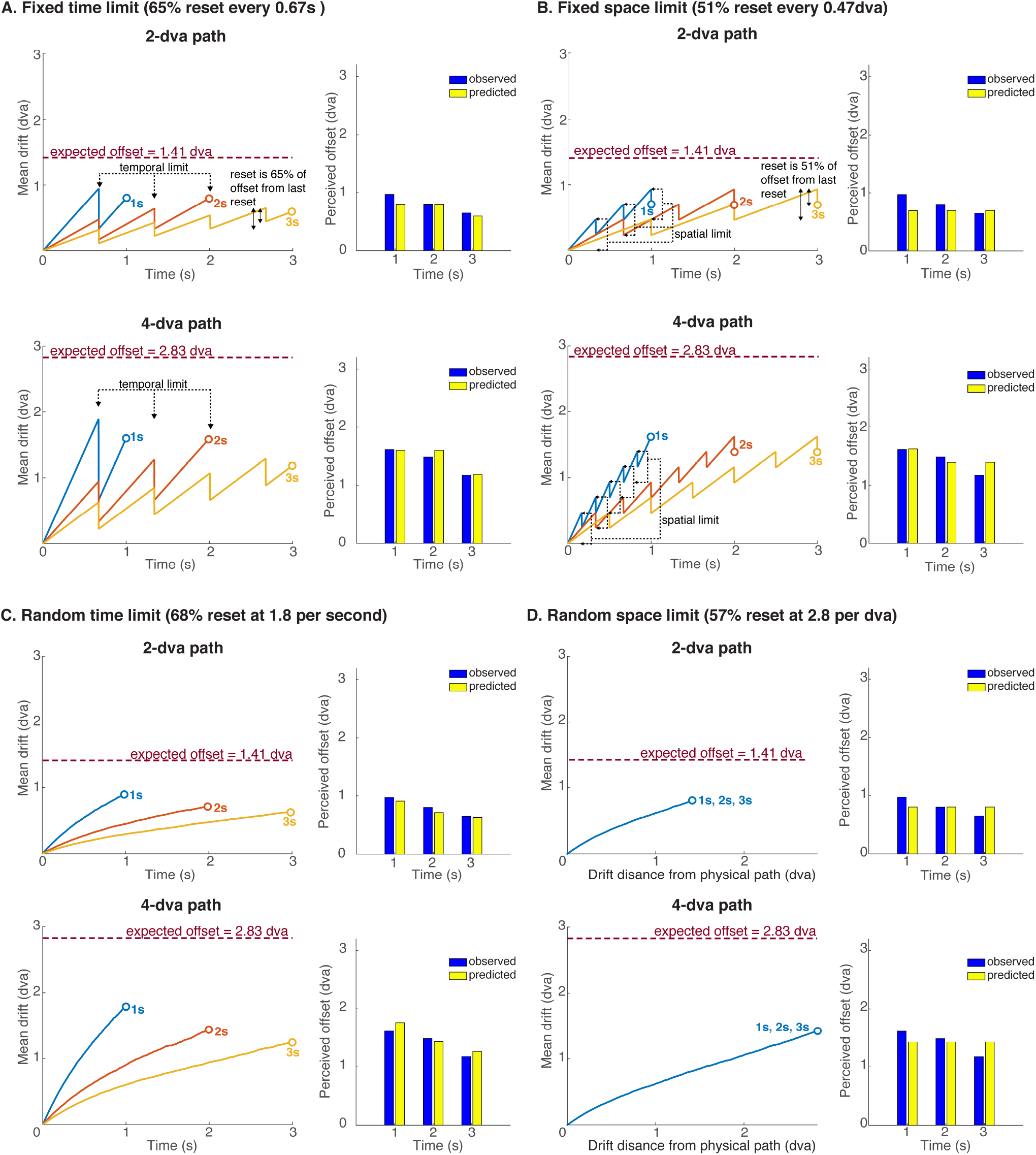
Partial reset model comparison. Average drift values across 10,000 simulations at each time point using the parameters that yielded lowest root mean square error (RMSE) between the observed PSEs and the predicted values (left panel) and the predicted and observed values at each path duration (1s, 2s and 3s) (right panel) for the A) fixed time limit model with 65% reset every 0.67s, B) fixed space limit model with 51% reset at 0.47 dva from last reset, C) random time limit model with 68% reset at 1.8/s and D) random space limit model with 57% reset at 2.8/dva.

**Table 1.** Mean number of resets for each path duration and length across 10,000 runs of simulation of the best fit models with full and partial resets and model performance.

| *Model* | Reset size<br>(% distance<br>to last reset) | Limit | Mean number of resets |  |  |  | AIC<sub>c</sub> | ΔAIC<sub>c</sub> |
| --- | --- | --- | --- | --- | --- | --- | --- | --- |
|  | Full resets<br>100% |  | Path<br>length | Path<br>duration |  |  |  |  |
|  |  |  |  | 1s | 2s | 3s |  |  |
| *Fixed time* |  | 1s | 2dva<br>4dva | 0<br>0 | 1<br>1 | 2<br>2 | -4.33 | 17.09 |
| *Fixed space* |  | 1.62dva | 2 dva<br>4 dva | 0<br>1 | 0<br>1 | 0<br>1 | -5.63 | 15.79 |
| *Random time* |  | 0.8/s | 2 dva<br>4 dva | 0.82<br>0.80 | 1.62<br>1.61 | 2.41<br>2.41 | -18.7 | 2.72 |
| *Random space* |  | 0.6/dva | 2 dva<br>4 dva | 0.84<br>1.70 | 0.84<br>1.70 | 0.84<br>1.70 | -16.3 | 5.12 |
|  | Partial resets |  |  |  |  |  |  |  |
| *Fixed time* | 65% | 0.67s | 2dva<br>4dva | 1<br>1 | 2<br>2 | 4<br>4 | -21.23 | 0.19 |
| *Fixed space* | 51% | 0.47dva | 2dva<br>4dva | 3<br>5 | 3<br>6 | 3<br>6 | -14.27 | 7.15 |
| *Random time* | 68% | 1.8/s | 2dva<br>4dva | 1.78<br>1.83 | 3.59<br>3.60 | 5.43<br>5.32 | -21.42 | 0 |
| *Random space* | 57% | 2.8/dva | 2dva<br>4dva | 3.94<br>7.88 | 3.94<br>7.88 | 3.94<br>7.88 | -13.91 | 7.51 |

We first examined models that assumed that spontaneous resets occurring prior to the temporal gap returned the perceived path 100% of the way to the physical location. For the fixed space and time model, with no random factor, the predicted offset just oscillates back and forth between 0 and the value set by the spatial or temporal limit (**Figure 10A** and **B).** These models with a fixed limit are not very realistic as the resets occur at the same point on every trial. Although observers often report seeing spontaneous resets (see ’t Hart, Henriques, & Cavanagh, 2019), they seem more variable. Nevertheless, the model predictions as shown in **Figure 10A** and **B** do demonstrate what an individual trace might look like with resets. Thus, the predicted PSE from a model with fixed time limit *t* would be a value between 0 and the expected offset after *t* seconds without a reset (as given in Equation 2; **Figure 10A**). Similarly, the prediction with a fixed spatial limit is just a value between 0 and the spatial limit *s* for both path lengths and all three durations (**Figure 10B**). Note that in these fixed models, the expected values of the perceived offset at each time point do not change across simulation runs. As shown in **Figure 10B,** a fixed space limit model with a spatial limit at 1.62 dva from the physical location is not a good match to the observed data as there is no difference in illusory offsets between the three path durations and the predicted offsets do not increase for the longer path length. On the other hand, the fixed time limit model does predict a decrease in the offset with duration. The rate of accumulation at each time point (Equation 2) is proportional to 1/duration so predicting the offsets at 1, 2 and 3s with ratios of 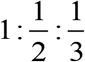. The observed data does approximately follow this trend i.e. decreasing offsets over time. However, the PSEs are overestimated for the 1s path as compared to the observed data (**Figure 10A** right panel). In addition, when the path length doubles for a fixed duration, the number of resets is unaffected because it is determined by time only, so the predicted offset doubles as well (directly proportional to the path length, Equation 2). The observed data do approximately double with the doubled path length (ratio of 4-dva to 2-dva of the observed PSE is 1.75).

The random models allow resets to occur probabilistically at some rate of resets per time interval or per spatial interval. As shown in **Figure 10C** and **D**, the average predictions of offset for these models increase smoothly due to the averaging over 10,000 runs of simulation with resets occurred randomly over time; each individual repetition had a sawtooth pattern of drift values like those in **Figure 10A** and **B.** Specifically, for the random time limit model with a fixed probability of reset over time, the rate that yielded the lowest RMSE was 0.8/s (on average one reset per 1.25s). As **Figure 10C** shows, similar to the fixed time limit model, predicted PSEs from this model decreased as path duration increased with fixed path length and doubled as the path length doubled. This pattern of results is expected from a model with random resets that are purely determined by the time the drift is away from the physical path since the number of resets will increase with time regardless of the path length (**Table 1**). With a fixed duration giving the same number of resets, doubling the path will double the PSE since the rate of drift increases directly with the external speed. However, unlike the fixed time limit model, the predicted PSE was smaller for 1s and the decrease in PSEs flattened over time for longer durations (**Figure 10C**; ratio of 1s to 2s to 3s of predicted PSEs is 1:0.7:0.5). The predicted PSEs from this model with a rate of one full reset per 1.25s matched the observed data better than the fixed time limit model (Δ*AIC_c_* = 14.37).

For the random space model (**Figure 10D**) with a constant probability over distance away from the physical path, the probability (*p*) that yielded the lowest RMSE was 0.6/dva (on average one reset per 1.7 dva). First, the predicted PSEs remained the same for the three path durations since the number of resets will stay the same as long as the drift distance is the same regardless of the path duration (**Table 1**). Also, unlike the case in the random time model, the predicted PSEs were on average less than doubled when the path length doubled (ratio of 4-dva to 2-dva of predicted PSEs is 1.4). This is because the number of resets doubles as the path length doubles, preventing the PSEs from doubling (**Table 1**). Thus, the predicted PSEs from this model at 0.6/dva did not fit the observed data as well as the random time model (Δ*AIC_c_* = 2.4).

Lastly, we examined the same types of fixed and random limit models with partial resets where the perceived path returned only part way to the last reset (**Figure 11** and **Table 1**). The pattern of results was similar to those with full resets. The best fit model had resets occurring randomly at a rate of 1.8/s (on average one reset per 0.56s) and each reset returned 68% of offset to last reset. Predictions from this model matched the data better than the fixed and random space models when they were allowed to have partial resets (Δ*AIC_c_* > 7). Predictions from the fixed time model with partial resets (65% partial resets every 0.67s) also matched the data as well as the random time model with partial resets (Δ*AIC_c_* = 0.19). As expected, the average number of resets of the best fit model with partial resets is higher than that with full resets to compensate for the smaller resets (**Table 1**).

Thus, the model that fits the observed PSEs best was the one where reset was partial (68%) and was set by random events at a rate of one every 0.56s (1.8 per second).

## Discussion

The present study examined the spatial and temporal limitations of the motion-position integration of the double-drift stimulus. By introducing a temporal gap in the middle of the motion path of a double-drift stimulus, we measured the magnitude of position accumulation up to a point for different path length and durations. Overall, we found that the perceived offset between the pre- and post-gap motion segments decreased as time increased and approximately doubled when the length of the motion path doubled. Our results are best characterized by a model in which a reset of the illusory drift occurs randomly with a constant rate over time. This random time model captures two main properties of the data. First, as the duration increases, the number of resets increases (linearly) and as a result, for a fixed path length, the perceived offset must decrease with duration. This matches the effect of duration in the data with path length fixed (**Figure 9**). In contrast, a purely spatial limit would predict the same illusory offset for a fixed path length independently of the duration of the path as the resets would be triggered purely by distance (since the illusory direction was titrated to be 45° for all stimulus conditions, the expected drift is set only by the path length and is therefore 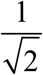 of path length). Second, with a purely temporal limit, the number of resets would stay the same for a given duration regardless of the path length. Doubling the path length would then double the illusory offset as the illusory drift accumulates at twice the rate when the path is twice as long (again, the expected drift is 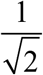 of two times of the path length), as observed in the data. A pure spatial limit model would predict more resets with increasing path length and that would keep the drift value from doubling. Thus, our results are best explained by a pure time limit where partial resets that return the perceived path 68% of offset to last reset that happens randomly, with a fixed rate across time on average once every 0.56s, independently of the magnitude of the Gabor’s perceived shift away from its physical location.

Although the fixed time model with partial resets also fit the data relatively well, models with a fixed limit are problematic both because the resets are not observed recurring reliably at the same position in individual trials (’t Hart et al., 2019) and because the model predictions themselves are very unstable – slight shifts of the spatial or temporal limits make very large changes in the predictions. Nevertheless, they do help us visualize the variability of the paths with position resets. These position resets, when occurring randomly over time, result in a more or less smooth trajectory that is curved across time. We also tested models that combined both space and time limits but already with only 6 data points, our current models stretch the ability of the data to constrain the models. With even more free parameters, the marginal gain with combined space and time limits is very small and they did not perform noticeably better.

Our results are best explained by resets occurring randomly over time with underlying linear illusory paths. These models predict curved paths when the position offsets are averaged over trials (**Figure 10 and 11**) and are quite similar to the curved paths described by Shapiro et al. (2010) and Kwon et al. (2015), although they predict that the perceived path is curved on each transit whereas we assume linear paths interspersed with resets. The data in our experiments here do not differentiate between these two alternatives. The demonstration movie from Lisi and Cavanagh (2015) shows a gap-triggered reset of linear path to almost all observers. Hand tracing data from ’t Hart, Henriques, and Cavanagh (2019) and Nakayama and Holcombe (2020) supports the existence of spontaneous resets along with an otherwise a linear path (see **Figure 4**). However, the resets, when they do occur, are not very salient and observers are often left with a feeling that the path shifted but without knowing when that happened, so more studies will be required to reach a clear conclusion.

Several previous psychophysical studies have examined the motion-position integration process of the double-drift stimulus. Tse and Hsieh (2006) first reported that the large deviation in perceived direction was produced by a combination of its local and global motion vectors. Lisi and Cavanagh (2015) further showed that the deviation is not only in direction but also in the perceived position of the stimulus, so that a brief temporal gap in the middle of the path would reset the illusory position offset back to its physical location. They proposed that the large and sustained perceived position displacement for the double-drift stimulus reflects the accumulation of local position shifts driven by the motion of its internal texture, shifting the perceived path far from the physical path (Lisi & Cavanagh, 2015). Note that in their study, the perceived offset between the pre- and post-gap segments was not significantly different from the expected offset (1.41 dva) for a 2dva motion path with 45° illusory angle (Lisi & Cavanagh, 2015), unlike our results in which we see a decrease in perceived offset for this condition (**Figure 9**). We assume that this difference was due to a difference in the speed of internal motion used in the two studies. In our study, we asked the participants to adjust the speed of the internal motion until the motion path appeared to be, locally, 45° away from its physical path for each external speed condition, ignoring any shifts due to spontaneous resets. These titrated internal motion speeds were all shown to be around the expected illusory angle (45°) derived from a vector sum model of the adjusted internal motion speed and the external motion speed (**Figure 6B**). Importantly, the internal motion speed that yielded a 45° illusory angle for a 2 dva/s motion path, which was then used in the subsequent reset test, was around 2 Hz in our study. However, Lisi and Cavanagh (2015) used a 3 Hz internal motion, which should yield a 56° illusory angle and an expected offset of 1.66 dva if there were no resets before the gap based on a vector sum model. Thus, if we assume that the perceived motion direction of the double-drift stimulus was produced by a vector sum of the external and internal motion speed when uncertainty is high as shown in Cavanagh and Tse (2019), it is possible that Lisi and Cavanagh (2015) also got less than the expected offset at the moment of the temporal gap.

The double-drift stimulus has a much longer and more sustained motion-position integration process than that of other well-known motion-induced position shift effects. For example, the “flash-grab” illusion shows integration period of only about 80 to 100 ms (Cavanagh & Anstis, 2013). The motion drag illusion (R. L. De Valois & De Valois, 1991; Eagleman & Sejnowski, 2007) shows an increase in the perceived offset that grows with presentation duration up to 50 or 60 ms (Kosovicheva et al, 2014; Jeon et al., 2020; Chung, Patel, Bedell, & Yilmaz, 2007). Unlike these other motion-induced position shifts, the double-drift stimulus involves a second-long accumulation of position errors. With such a long motion-integration period, it is unlikely that cells in the early visual areas with their short integration time windows are responsible for building up the position errors underlying this illusion. A recent neuroimaging study found that higher-order cortical areas such as the frontal cortex have a shared encoding of the illusory and matched physical path of the double-drift stimulus, whereas common representations were not observed in the early visual cortex (Liu et al., 2019). They proposed that while extrastriate areas could be encoding the local position errors driven by the combined internal and external motion vectors integrated over short durations, the higher-order brain areas are necessary to accumulate the outputs from these early regions and transform them into the final percept of the motion path (Liu et al., 2019). Importantly, the higher-order areas found in their study were mostly in anterior brain regions that were known to be involved in executive control and working-memory-related processing, such as the medial prefrontal cortex (MPFC) and lateral prefrontal cortex (LPFC). Thus, it would seem that the local position offsets computed in early visual cortices are fed into these working-memory-related frontal regions that act as ‘visual buffers’ where the position errors are accumulated and maintained over longer durations.

Moreover, studies of saccades and smooth pursuit to targets with motion induced position shifts also suggest a clear difference in time constants between the double-drift stimulus and other motion induced position shifts. First of all, position shifts along the motion path such as the flash-grab, motion drag, and flash-drag illusions show similar responses for saccades and perception (de’Sperati & Baud-Bovy, 2008; Schafer & Moore, 2007; Kosovicheva et al., 2014; van Heusden, Rolfs, Cavanagh, & Hogendoorn, 2018; Zimmermann, Morrone, & Burr, 2012). These motion-induced position shifts have short motion integration periods that are similar to the time over which the saccade system integrates motion and position signals to program a saccade to a moving stimulus (saccadic dead time, Becker 1991). This shared integration time for the saccade system and the perceptual system in the extrapolation of moving targets may keep perception and saccades aligned when acting on a moving stimulus. In contrast, as shown here and in Lisi and Cavanagh (2015), the double-drift accumulates a deviation orthogonal to the motion path for over a second, well beyond the duration over which motion integration operates for saccades. Moreover, it was shown that saccades appear to be immune to the double-drift illusion as eye movements are directed to the physical not the perceived location (Lisi & Cavanagh, 2015; although see Nakayama & Holcombe, 2020). Thus, it is possible that cortical systems with short information integration times such as the saccade system and iconic memory compute target positions in retinotopic coordinates and are not part of the network that computes the perception of the second-long deviation of the double-drift stimulus. This mismatch of temporal integration underscores the dissociation of position information for saccades and perception (Lisi & Cavanagh, 2015). Moreover, Cavanagh and Tse (2019) found that the illusion persists during smooth pursuit of a fixation spot that moves in tandem with a double-drift stimulus in the periphery, keeping it relatively fixed on the retina. This suggests that the computation of the illusion occurs at or beyond regions that compensate for eye movements to recover object locations in world coordinates.

Our study further explored the possible spatial and/or temporal limits to the accumulation of position offsets for the double-drift stimulus. We found that a purely temporal limit best captured the properties of the data. In particular, our modeling suggested spontaneous partial resets of the double-drift stimulus occur randomly, on average once every 0.56s (a rate of 1.8/s), independently of the distance traveled, where each reset returned 68% of offset to last reset. These random resets with a fixed rate across time would produce the two main effects observed in the data: the decrease in illusory offset with duration and the doubling of illusory offsets with doubling of the path length. Thus, these results imply that there are occasional, random interruptions of accumulation triggered by external events that possibly distract attention. A recent study from Nakayama and Holcombe (2020) proposed that attentional shifts are a possible source for triggering full resets of the perceived path for this stimulus. Interestingly, these random resets bring the perceived location back to the target’s physical position either fully or partially – indicating that some part of the visual system must be keeping track of the physical location even as the perceived location moves farther and farther away from its retinal location. This may be the saccade system – Lisi and Cavanagh (2015) showed that eye movements to the double-drift target land as accurately as those to a control with no internal motion, indicating that the saccade system retains an accurate representation of the physical location of the double-drift stimulus. Alternatively, the reset location may just be the center of the very uncertain location information. Despite the uncertainty, the location distribution does have a centroid, and this may be what replaces the offset location when a reset happens. If this were the case, the reset location should be highly variable, whereas if the reset location was obtained from the saccade system, it should not be especially noisy.

We note that the rate of microsaccades, at greater than once per second (Kagan, Gur, & Snodderly, 2008; Martinez-Conde, Macknik, & Hubel, 2004; Tse, Baumgartner, & Greenlee, 2010), is comparable to the rate of resets posited here (1.8/s), which also falls within the high end of the range of the rate of spontaneous eyeblinks (Colzato, van den Wildenberg, van Wouwe, Pannebakker, & Hommel, 2009). Future work will examine whether microsaccades or eyeblinks trigger the positional resets examined here. For the moment, the results here point to some random events that trigger partial resets of the accumulation of position errors for the double-drift stimulus at a fixed rate over time, independent of its distance from its physical path.

## Conclusions

The double-drift illusion is formed by the accumulation of motion-induced position errors orthogonal to the physical motion path over longer than a second. Studies have shown that the perceived path of this illusion can reset back to its physical location by a temporal gap (Lisi and Cavanagh, 2015) or by attentional distracting events such as transient flashes presented around the stimulus (Nakayama & Holcombe, 2020). Here we examined if such position resets can happen spontaneously, and if so, whether they depend on the time or/and the distance traveled by the double-drift stimulus. By introducing a temporal gap, we measured the size of the accumulated illusory offset up to the gap for different path lengths and durations. We found that the perceived offset at the gap was smaller than the expected offset size if no reset had occurred before the gap, suggesting that spontaneous resets had occurred before the gap. In addition, the offset size decreased for longer durations at the same path length and approximately doubled when the path length doubled. The observed data are best explained by resets that are determined purely by the time since the perceived path drifted away from the physical path, regardless of the distance traveled, with the partial resets occurring randomly at a constant rate over time (on average once every half second).

## Acknowledgments

This material is based upon work funded in part by National Science Foundation Award (1632738), the Department of Psychological and Brain Sciences of Dartmouth College, and NSERC Canada.

